# NanoTag - an IgG-free method for mapping DNA-protein interactions

**DOI:** 10.1101/2024.07.12.603224

**Authors:** Maria A. Dimitriu, Rodrigo G. Arzate-Mejía, Leonard C. Steg, Pierre-Luc Germain, Isabelle M. Mansuy

## Abstract

Genome-wide profiling of DNA-protein interactions in cells can provide important information about mechanisms of gene regulation. Most current methods for genome-wide profiling of DNA-bound proteins, such as ChIP-seq and CUT&Tag, use conventional IgG antibodies to bind target protein(s), which limits their applicability to targets for which high affinity and high specificity antibodies are available. Here we describe NanoTag, a novel method derived from CUT&Tag that is IgG-free and uses a nanobody to profile DNA-protein interactions. NanoTag is based on an anti-GFP nanobody-Tn5 transposase fusion that allows mapping GFP-tagged proteins associated with chromatin in a fast and cost-effective manner. We demonstrate the utility of NanoTag by profiling the histone mark H3K4me3 via its binding partner TATA box-binding protein-associated factor 3 (TAF3) and the transcription factors Nanog and CTCF in mouse embryonic stem cells expressing GFP-tagged targets. For the targets examined, NanoTag data shows high correlation to CUT&Tag data and displays a similarly high signal-to-noise ratio. Overall, NanoTag provides a flexible, IgG-free and cost-effective method to generate high resolution DNA-binding profiles in cells or tissues.

## Background

Interactions between proteins and DNA are essential for genome organization and regulation. To identify these interactions, it is necessary to precisely map the binding sites of regulatory proteins on chromatin. Chromatin immunoprecipitation with sequencing (ChIP-seq) has long been the gold standard method for genome-wide mapping of histone modifications and transcription factor (TF) binding sites[1]. Recent alternative techniques, such as Cleavage Under Targets and Tagmentation (CUT&Tag)[2] and derivatives,[3–10] have been developed to allow *in situ* profiling of chromatin components in low-input unfixed samples using a fusion protein between the IgG-binding protein A and the Tn5 transposase. By profiling native targets, these techniques remove the need for cells crosslinking and provide more biologically relevant profiles of the regulatory landscape while avoiding the risk of epitope masking[2,11]. They allow to map binding sites in the genome at high signal-to-noise ratio and resolution, and require lower sequencing depth than ChIP-seq[12]. However, most of these methods require the use of IgG antibodies to bind the desired target loci and specific IgG are only available for a subset of regulatory proteins[13]. They also have a time-consuming and expensive production process. A common strategy to circumvent the lack of IgG availability against many targets is to tag the protein of interest with an epitope such as green fluorescent protein (GFP,) and use tag-specific antibodies[13].

We developed a method called NanoTag that uses nanobodies instead of IgG to identify protein binding sites on DNA. Nanobodies are small compact antibodies with high antigen binding affinity, high cell penetrability and no cytotoxicity that can be produced at a large scale and low cost in bacteria, thus eliminating the use of animals[14]. Unlike CUT&Tag, which requires multiple successive steps to recruit protein A-Tn5 to desired targets, NanoTag is a single step method that uses a fused anti-GFP nanobody-Tn5 protein directly binding to target proteins. We established NanoTag in mouse embryonic stem cells (mESC) and implemented it to profile the histone posttranslational modification H3K4me3 via its protein binder TAF3[15] and the transcriptional regulators Nanog and CTCF via GFP-tagged TAF3, Nanog and CTCF in transgenic cells. With NanoTag, we achieved highly reproducible binding profiles at high resolution and with low background noise for all targets, that correlated well with data obtained with CUT&Tag in the same cells. NanoTag is a highly versatile and cost-effective technique that allows low-input chromatin profiling using a short and straightforward protocol, without IgG of animal origin. We anticipate that NanoTag will be useful for high-throughput analyses of DNA binding proteins or epigenetic marks for which high-affinity IgG are not available, by using existing cell lines that express GFP-tagged targets.

## Results and discussion

### Indirect profiling of H3K4me3 sites in mESC

To benchmark our NanoTag method, we characterized the binding profile of the H3K4me3-binding protein TAF3 and the TF Nanog and CTCF in mESC expressing GFP-tagged versions of each protein (Fig. 1a,b and Fig. 2a,h, see Methods). We compared the quality of NanoTag binding profiles to profiles obtained by CUT&Tag on the same cell lines using an anti-GFP antibody. As negative control, we conducted NanoTag and CUT&Tag on wild-type (WT) mESC not expressing any GFP-tagged target. We chose CUT&Tag to benchmark NanoTag because it is technically the closest even if it is not the most commonly used method to profile TF. Like NanoTag, CUT&Tag also has the advantage over methods such as CUT&RUN and ChIP-seq to provide high quality binding profiles with less input material, fewer sequencing reads [2].

**Figure 1.**
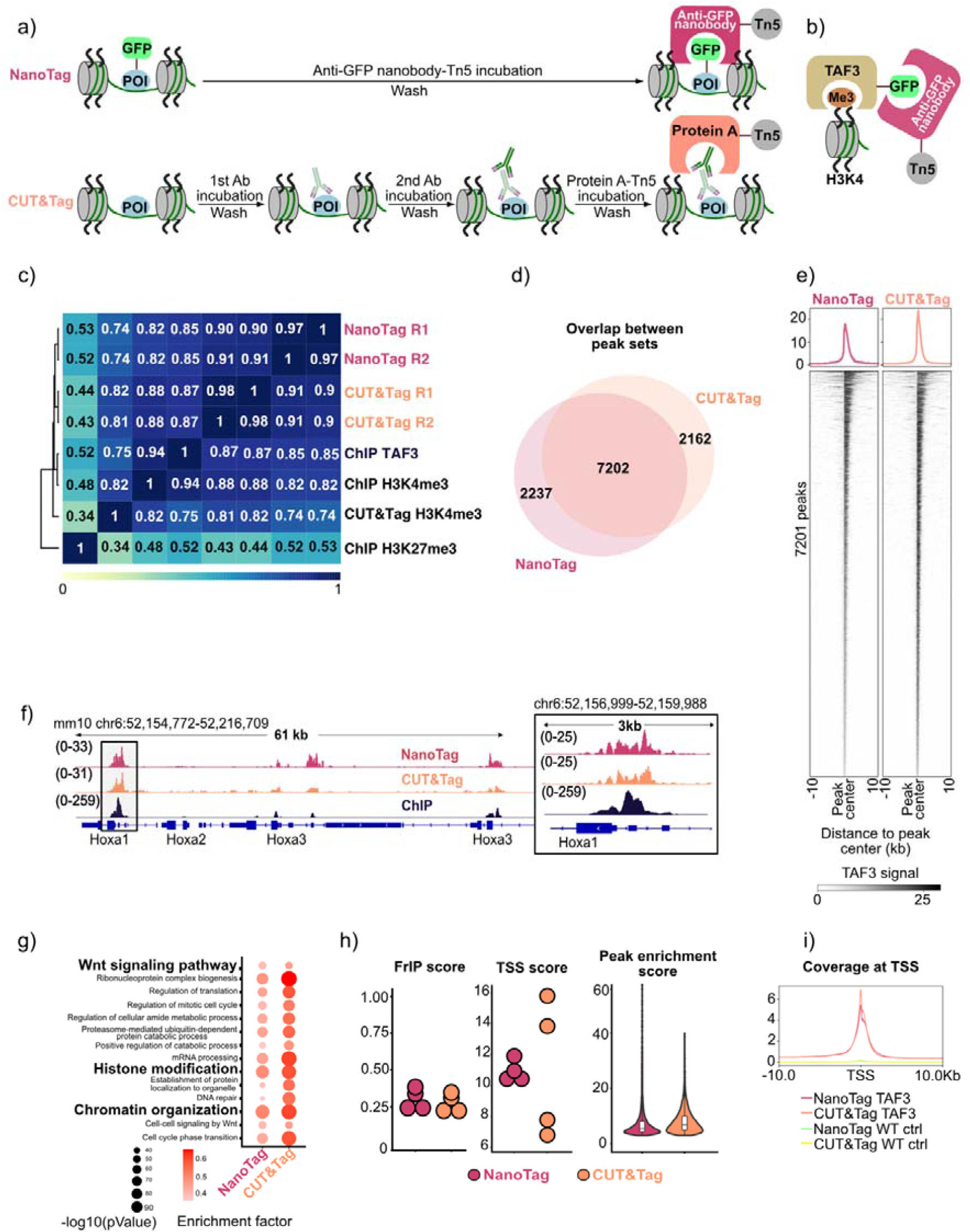
Comparison of NanoTag and CUT&Tag workflow and profiling of TAF3 binding. a) Schematic of the steps involved in NanoTag and CUT&Tag protocols. For NanoTag (magenta, top), permeabilized cells expressing a GFP-tagged protein of interest (POI) are incubated with an anti-GFP nanobody-Tn5 fusion protein. For CUT&Tag (orange, bottom) successive incubations with a primary antibody, a secondary antibody and protein A-Tn5 fusion are required. b) Schematic of GFP-tagged TAF3 protein expressed in mESC bound to H3K4me3 on the genome and detected by the anti-GFP nanobody-Tn5 with NanoTag. c) Hierarchically clustered correlation matrix of TAF3-NanoTag replicates (magenta, R1, R2), TAF3-CUT&Tag (orange, R1, R2), ChIP-seq targeting TAF3, H3K4me3 and H3K27me3 and CUT&Tag targeting H3K4me3. Spearman correlations were calculated using read coverage at TAF3 peaks common to NanoTag, CUT&Tag and ChIP-seq TAF3 datasets. NanoTag and CUT&Tag datasets were produced in-house, ChIP-seq TAF3 data are from[25], ChIP-seq H3K4me3 and H3K27me3 data are from[36] and CUT&Tag H3K4me3 data are from [35]. d) Overlap between NanoTag and CUT&Tag peak sets when profiling TAF3. The peak set corresponding to each method represents the union of peaks called in the two replicates. e) Profile plots and heatmaps showing TAF3-NanoTag (magenta) and TAF3-CUT&Tag (orange) enrichment around the 7201 common peaks. f) Genome browser view showing coverage of TAF3 NanoTag, TAF3-CUT&Tag and H3K4me3-ChIP-seq around the *Hoxa* locus. Close-up shows coverage of the same data around *Hoxa1*. g) Top 14 common GO terms identified as enriched in TAF3-NanoTag and TAF3-CUT&Tag peak sets. h) Plots summarizing the FrIP scores (calculated as fraction of reads from each dataset falling into ChIP-seq TAF3 peaks), TSS scores and peak enrichment score distribution for NanoTag and CUT&Tag TAF3 data. FrIP and TSS scores were calculated individually for peak sets called in each replicate. Coverage of individual replicates was merged to calculate the enrichment score at common peaks. i) Profile plot comparison of TAF3-NanoTag and CUT&Tag signal around TSS in TAF3-GFP and wild-type (WT) mESC controls. For each method, the shown profile represents the merged coverage of two replicates.

**Figure 2.**
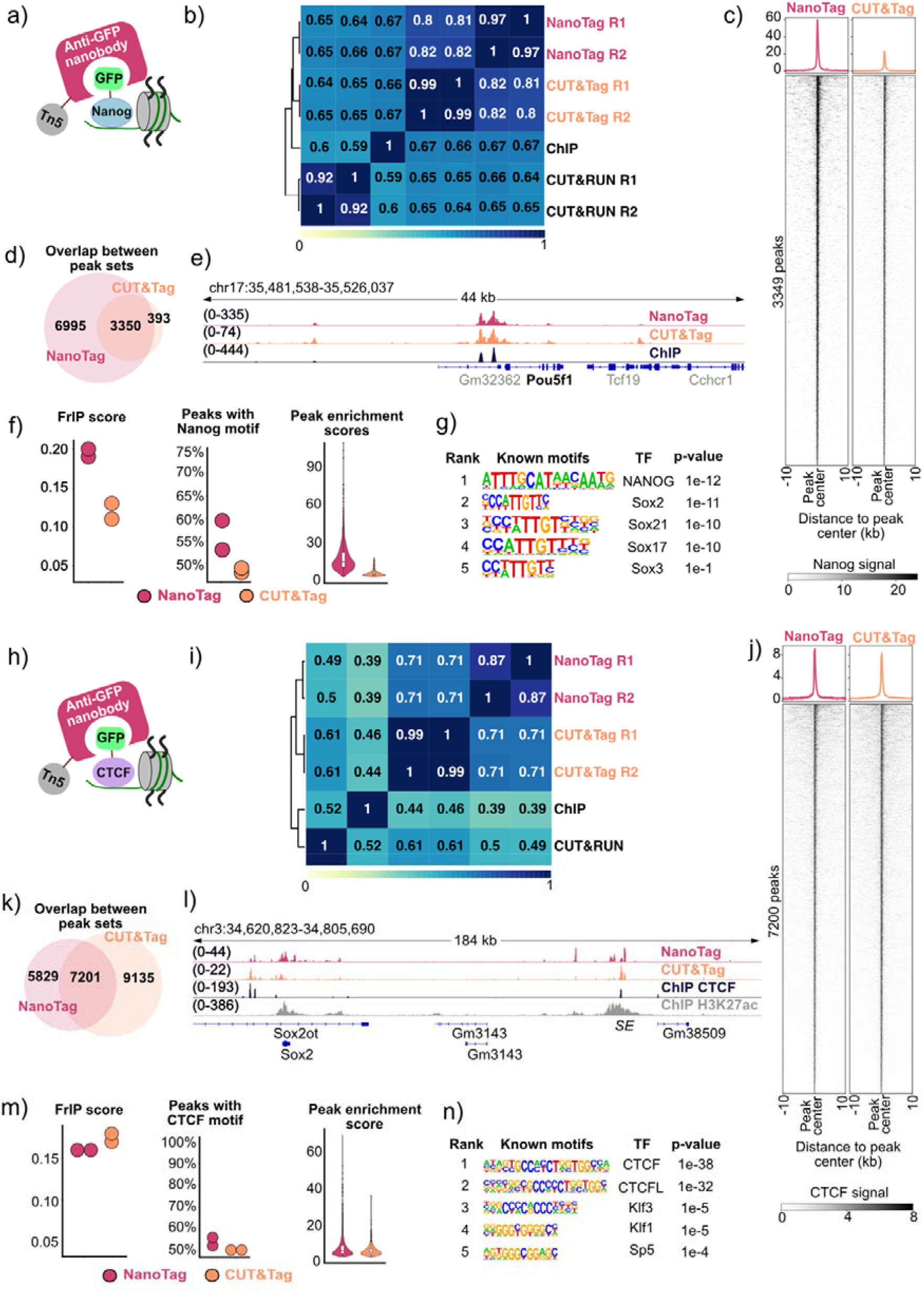
Nanog and CTCF binding sites captured by NanoTag. a, h) Schematic of GFP-tagged Nanog (a) or CTCF (h) expressed in mESC bound to DNA and detected by the anti-GFP nanobody-Tn5 with NanoTag. b) Hierarchically clustered correlation matrix of NanoTag replicates (magenta, R1, R2), CUT&Tag (orange, R1, R2), ChIP-seq and CUT&RUN (R1, R2) data targeting Nanog. Spearman correlations were calculated using read coverage at Nanog peaks commonly called in NanoTag, CUT&Tag and ChIP-seq datasets. NanoTag and CUT&Tag datasets were produced in-house, ChIP-seq data are from[36] and CUT&RUN data are from[35]. c) Profile plots and heatmaps describing Nanog-NanoTag (magenta) and Nanog-CUT&Tag (orange) enrichment around the 3349 common peaks. For each method, the signal was merged across two replicates. d) Overlap between NanoTag and CUT&Tag peak sets when profiling Nanog data. The peak set corresponding to each method represents the union of peaks called in the two replicates. e) Genome browser snapshot showing NanoTag, CUT&Tag and ChIP-seq profiles targeting Nanog around the *Pou5f1* locus. The profile for each method represents the signal obtained from merging two replicates. f) Plots summarizing FrIP scores at ENCODE peaks, the proportion of peaks containing Nanog motifs and the enrichment score distribution for common peaks when profiling Nanog using NanoTag (magenta) and CUT&Tag (orange). FrIP scores and proportion of peaks containing the Nanog motif were calculated individually for peak sets called in each replicate. Coverage of individual replicates was merged to calculate the enrichment score at common peaks. g) Most highly enriched TF motifs in NanoTag peaks when profiling Nanog. Motifs were identified in the 500 most significant peaks across the two replicates. h) Same as a) but profiling CTCF. i) Hierarchically clustered correlation matrix of NanoTag replicates (magenta, R1, R2), CUT&Tag (orange, R1, R2), ChIP-seq and CUT&RUN data targeting CTCF. Spearman correlations were calculated using read coverage at CTCF peaks commonly called in the NanoTag, CUT&Tag and ChIP-seq datasets. NanoTag and CUT&Tag datasets were produced in-house, ChIP-seq data are from[36], CUT&RUN data are from[34]. j) Same as c) but profiling CTCF. k) Same as d) but profiling CTCF. l) Genome browser snapshot showing NanoTag, CUT&Tag and ChIP-seq data profiling CTCF and ChIP-seq profiling H3K27ac at *Sox2* locus. NanoTag and CUT&Tag data were generated in-house, CTCF ChIP-seq data are from[36], H3K27ac ChIP-seq data are from[43]. SE = *Sox2* superenhancer. m) Same as f) but profiling CTCF. n) Same as g) but profiling CTCF.

NanoTag and CUT&Tag experiments were performed in duplicate using 400,000 cells per sample and libraries were prepared in parallel (see Methods for NanoTag, CUT&Tag). Before sequencing, we confirmed the enrichment of NanoTag libraries for regions bound by the desired protein by qPCR. As expected, we observed that libraries produced from cells expressing the GFP-tagged target showed up to 15,000-fold faster amplification (target-dependent) at target regions compared to WT libraries (Fig. S1).

When profiling TAF3 (TAF3-NanoTag), NanoTag coverage showed high reproducibility across replicates and correlated well with both TAF3-CUT&Tag and H3K4me3-CUT&Tag peak data (91% and 74% respectively) (Fig. 1c). TAF3-NanoTag coverage at peaks showed high correlation with H3K4me3 but not with H3K27me ChIP-seq in mESC (Fig. 1c). Importantly, the majority of peaks called in the TAF3-NanoTag data (76%) overlapped with those called in CUT&Tag data (Fig. 1d) and common peaks showed similar signal coverage (Fig. 1e). TAF3-NanoTag peaks were located predominantly at promoters (Fig. S2), consistent with TAF3 binding to H3K4me3, a histone modification enriched at promoters. We observed that NanoTag coverage is similar to that of CUT&Tag at *Hox* genes cluster, where H3K4me3 signal is expected in mESC[16,17] (Fig. 1f). TAF3-NanoTag peaks were also found to be enriched at genes associated with chromatin organization and histone modification (Fig. 1g). TAF3-NanoTag demonstrated comparable scores for fragments in peaks (FrIP), transcription start site (TSS) and peak enrichment when compared with TAF3-CUT&Tag (Fig. 1h), indicating that NanoTag is similar in sensitivity to CUT&Tag when profiling TAF3. In contrast, sparse landscapes were produced by both methods in control cells (WT), suggesting comparably low background noise (Fig. 1i). We observed a higher overlap of NanoTag and CUT&Tag TAF3 peaks with promoter regions than for ChIP-seq (Fig.S2), which is consistent with the previously reported bias for Tn5-based methods toward genomic regions with more open chromatin[18]. Together, these results demonstrate the suitability of NanoTag to profile H3K4me3 sites in the genome of mESC using one of its binding partners.

### Direct profiling of Nanog and CTCF binding sites in mESC

Next, we used NanoTag to profile the binding of the TF Nanog and CTCF in mESC expressing GFP-tagged Nanog[19] and GFP-tagged CTCF[20] (Fig. 2a,h). NanoTag data showed high reproducibility across replicates when profiling Nanog (Fig. 2b) and CTCF (Fig. 2i), high correlation with CUT&Tag and lower correlation with CUT&RUN and ChIP-seq data targeting the respective TF (Fig. 2b,i). For Nanog, NanoTag showed higher coverage at peaks common to NanoTag and CUT&Tag (Fig. 2c) and at ENCODE Nanog peaks (Fig. S3a). For CTCF, NanoTag and CUT&Tag coverage was similar at common peaks (Fig. 2j) and at ENCODE CTCF peaks (Fig. S3a). NanoTag captured 90% of CUT&Tag peaks when profiling Nanog (Fig. 2d) and 44% of CUT&Tag peaks when profiling CTCF (Fig. 2k). However, NanoTag data showed coverage also at peaks uniquely called for CUT&Tag for all targets, including CTCF (Fig. S4), indicating that the low peak overlap between techniques may be influenced by the peak-calling strategy. Low peak overlap has been reported for various targets across replicates using a single technique e.g. CUT&Run [21]. We observed similar signal for both methods at *Pou5f1* locus when profiling Nanog (Fig. 2e) and at *Sox2* locus when profiling CTCF (Fig. 2l). NanoTag showed higher FrIP scores, more peaks containing the TF motif and higher peak enrichment scores compared with CUT&Tag when profiling Nanog (Fig. 2f), whereas similar scores were found when profiling CTCF (Fig. 2m). Further, in line with expectations, NanoTag peaks showed the highest enrichment in Nanog and CTCF motif when profiling the respective TF (Fig. 2i, n). 70% of the top 50 gene ontology (GO) terms overlapped for NanoTag and CUT&Tag when profiling Nanog, and 72% when profiling CTCF. Wnt signaling, a key regulator of mESC self-renewal, also appeared among the top GO terms in the NanoTag data (Fig. S5). Finally, NanoTag identified fewer peaks than CUT&RUN and ChIP-seq when profiling TF (Fig. S6), which may partially be due to lower sequencing depth used for NanoTag (Table S1,S2). These results confirm that NanoTag can be used to directly profile Nanog and CTCF binding sites in mESC.

### Versatility of NanoTag

We tested the applicability of NanoTag to different input material by targeting TAF3 in fresh mESC, and in fresh or frozen nuclei from mESC (see Methods). We obtained similar profiling results across conditions (Fig. S7), suggesting that freezing does not reduce NanoTag library complexity. Since concanavalin A beads used to bind cells or nuclei are expensive, we also tested a version of NanoTag protocol for which cells/nuclei are centrifuged during collection and washing steps instead of bound by beads (see Methods). This produced highly similar results across protocol versions and inputs (Fig. S7), suggesting high versatility of NanoTag.

### Limitations

One shortcoming of NanoTag is that it has a high rate of duplicated reads, an issue also reported for CUT&Tag[22] (Table S1). However, this did not alter the quality of the results since very few reads were sufficient to obtain high quality binding profiles. Further, if needed, this can be circumvented by sequencing more deeply to obtain enough unique DNA fragments despite high duplication rates, as sequencing has become reasonable cost-wise[23]. In its current form, NanoTag requires cells expressing GFP-tagged targets but such cells are widely available or can be easily generated using CRISPR technologies[24]. We expect NanoTag to be broadly applicable to various chromatin targets but we can so far draw conclusions only for targets we tested. We are planning to test additional targets in the future, including profiling patterns of methylated CpGs by applying NanoTag to MBD1-GFP expressing mESC[25]. Further, one derivation of NanoTag we envision, amenable to *in vivo* applications, is to use fusions between Tn5 and nanobodies or other antibody mimetics that directly bind an endogenous target. Such binders of chromatin-associated proteins already exist[26–29] and are continuously being developed against new targets.

## Conclusions

We successfully developed NanoTag, a novel method to map binding sites of chromatin-associated proteins and applied it to profile the genomic location of H3K4me3, Nanog and CTCF in mESC. NanoTag provides high reproducibility and signal-to-noise ratio and delivers signals similar to CUT&Tag. NanoTag offers a significant advantage over other methods by being IgG-free, making it suitable for profiling also regulatory proteins that lack high-quality antibodies[13]. Additionally, NanoTag is a highly cost-effective technique. It does not depend on expensive antibodies that involve extensive animal use to be produced. Instead, it utilizes a nanobody-Tn5 fusion that can be produced inexpensively and in large quantities using bacteria. It is a versatile and efficient method for high-throughput characterization of DNA binding expected to be an attractive alternative for epigenomic profiling across cells and tissues.

## Methods

### Transposome preparation

Using the 3XFlag-pA-Tn5-Fl expression vector[2] (Addgene plasmid #124601), protein A sequence was replaced with an anti-GFP nanobody sequence that was PCR amplified from the pGEX6P1-GFP-Nanobody vector [30] (Addgene plasmid #61838) to add restriction sites for *HindIII* and *EcoRI* flanking the anti-GFP nanobody sequence. The anti-GFP nanobody PCR product and the 3XFlag-pA-Tn5-Fl vector were digested with *HindIII* and *EcoRI* for 1 hour at 37^°^C. The digested PCR product was purified using the QIAquick Gel Extraction Kit (Qiagen, cat.no. 28115) and the digested vector was purified on a 2% agarose gel. Digestion products were ligated for 1 hour at 22^°^C then transformed into DH5α competent bacteria. Colonies were picked and sequenced to verify the sequence of the anti-GFP nanobody-Tn5 (pMD1) plasmid (Addgene #215567). The anti-GFP nanobody-Tn5 protein fusion was produced by the Protein Production and Structure Core Facility of the Swiss Federal Institute of Technology (EPFL), Lausanne, Switzerland, following the protocol described by Chen et al. for Tn5 purification[31]. Briefly, *E.coli* LysY/Iq competent bacteria were transformed with the pMD1 plasmid. One colony was used to inoculate 2L of LB bacterial growth media and overexpression of the anti-GFP nanobody-Tn5 was induced with 0.25 mM IPTG at 18^°^C overnight. The bacterial pellet was resuspended in HEGX buffer (NaCl 800mM, HEPES 20mM pH 7.5, EDTA 1mM, Triton X-100 0.1%, Glycerol 10% + protease inhibitor cocktail), sonicated for 2.5 min using 10 second cycles at 70% amplitude and centrifuged for 30 min at 20,000xg. To remove bacterial DNA, the supernatant was treated with 0.275% PEI and centrifuged for 20 min at 20,000xg. The supernatant was bound to chitin beads for 2h at 4^°^C, washed with HEGX buffer and cleavage of the anti-GFP nanobody-Tn5 from the intein tag was induced with 100 mM DTT overnight, eluted and loaded with oligonucleotides for 1 hour at 30^°^C. Oligonucleotides (synthesized by IDT) were resuspended at 200 μM in water, heated to 95^°^C for 5 min and slowly cooled down to hybridize, then mixed and heated to 95^°^C for 15 min then cooled down to 50^°^C and diluted 1:1 in 2xHEGX buffer with no Triton-X. The protein was dialysed overnight, concentrated, diluted 1:1 in 50 mM HEPES, pH 7.5, 100 mM NaCl, 0.1 mM EDTA, 1 mM DTT, 50% Glycerol, aliquoted, flash-frozen and stored at –80^°^C. 2L of bacterial culture yielded 22.5 mL at 23 μM of anti-GFP nanobody-Tn5 transposomes.

### Oligonucleotides

FWD primer for anti-GFP nanobody containing *HindIII* site: GATCAAGCTTAC

REV primer for anti-GFP nanobody containing *EcoRI* site: CAGAATTCGATC

Tn5-A: TCGTCGGCAGCGTCAGATGTGTATAAGAGACAG

Tn5-B: GTCTCGTGGGCTCGGAGATGTGTATAAGAGACAG

Tn5-rev: /5Phos/CTGTCTCTTATACACATCT

### Cell lines and culture

Wild-type mESC (HA36CB1, 129×C57BL/6) and mESC expressing GFP-tagged engineered chromatin readers (TAF3 for H3K4me3)[25] were kindly provided by Tuncay Baubec. R1 mESC expressing Venus-tagged Nanog[19] were kindly provided by Timm Schroeder. E14Tg2a AID-eGFP-CTCF, Tir1 mESC[20] were kindly provided by Elphège Nora. Of note, CTCF expression levels are 2-3-fold lower in the AID-eGFP-CTCF compared to wild-type cells[20]. mESC were cultured on 0.2% gelatin-coated culture dishes in Knockout DMEM supplemented with 15% Fetal Bovine Serum (FBS) qualified for ES cells, 1X GlutaMax, 1X Penicillin/Streptomycin, 1X MEM Non-Essential Amino Acid, 100 μM β– Mercaptoethanol, and 1000 U/ml Leukemia inhibitory factor. Media was changed daily and mESC were passaged every 2 days.

### NanoTag

A detailed step-by-step protocol can be found on protocols.io. All NanoTag experiments were conducted in duplicate. For all experiments, 400,000 fresh mESC were collected by treatment with StemPro Accutase at 37^°^C for 5 min and centrifugation at 600xg for 3 min and washed with Wash buffer (20 mM HEPES pH 7.5, 150 mM NaCl, 0.5 mM spermidine, protease inhibitors). Concanavalin A (conA) beads were washed with Binding buffer (20 mM HEPES-KOH pH 7.9, 10 mM KCl, 1 mM CaCl_2_, 1 mM MnCl_2_) as described for CUT&Tag[32]. 10 μL of conA bead slurry was added to the 400,000 cells while gently vortexing and incubated on a rotator for 15 minutes. Cells bound to conA beads were then resuspended in 100 μL of Dig-wash buffer containing 2 mM EDTA and 2 μL of anti-GFP nanobody-Tn5 transposomes and incubated overnight on a nutator at 4^°^C. Cells were washed 3 times with 1 mL of Dig-300 buffer by incubating on a nutator for 5 min, then resuspended in 300 μL Dig-300 buffer with 10 mM MgCl_2_ and incubated at 37^°^C for 1 hour. Tagmentation was stopped and DNA was extracted using phenol-chloroform extraction as described for CUT&Tag [32]. When performing NanoTag without the use of conA beads, cells were instead centrifuged at 600xg for 5 min at 4^°^C for every resuspension step. When performing NanoTag on nuclei, cells were treated with Nuclei extraction buffer (20 mM HEPES-KOH pH 7.9, 10 mM KCl, 0.5 mM Spermidine, 0.1% Triton X-100, 20% glycerol, protease inhibitors), as described[32]. When performing NanoTag on frozen nuclei, cells were treated with Nuclei extraction buffer and nuclei were frozen as described[32]. Libraries were prepared simultaneously for NanoTag and CUT&Tag samples. To determine the amount of cycles necessary for each library, a 2.5 µL aliquot of the tagmented DNA was amplified by qPCR using 2 µL of i5 index, 2 µL of i7 index, 7.5 µL of NEBNext PCR Master mix, 0.15 µL of 100x SybrGreenI dye and 0.85 µL of water with the following program: 72^°^C for 5 min, 98^°^C for 30 seconds, and 30 cycles of: 98^°^C for 10s, 63^°^C for 30s and 72^°^C for 1 min[32]. 21 µL of tagmented DNA was then PCR amplified with the amount of cycles determined by qPCR to yield one third of the maximum fluorescence using 2 µL of i5 index, 2 µL of i7 index and 25 µL of NEBNext PCR Master mix as described for ATAC-seq [33]. NanoTag libraries were amplified using 21-22 cycles by PCR (72^°^C for 5 min, 98^°^C for 30 sec, n cycles of 98^°^C for 10 seconds and 63^°^C for 10 seconds, 72^°^C for 1 min and held at 8^°^C) to achieve a library concentration of at least 2 nM. Libraries were cleaned up as described[32] using 60 µL of AMPure beads and eluted in 20 µL of 10 mM Tris-HCl pH 8.0. The library profile was verified by Tapestation with the High-sensitivity D1000 kit.

### CUT&Tag

CUT&Tag experiments were conducted in duplicate as described[32] with minor modifications. For all experiments, 400,000 fresh mESC were collected, washed with Wash buffer and bound to conA beads as described above for NanoTag. Beads were then resuspended in 50 μL Dig-wash buffer (20 mM HEPES pH 7.5, 150 mM NaCl, 0.5 mM spermidine, protease inhibitors, 0.05% digitonin) with 0.1% BSA, 2 mM EDTA and 1:100 anti-GFP antibody (Abcam, cat. no. ab290) or IgG (EpiCypher rabbit IgG, cat. no. 13-0042) and incubated overnight on a nutator at 4^°^C. Beads were resuspended in 100 μL of Dig-wash buffer with 1:100 secondary antibody (guinea pig α-rabbit antibody (Antibodies online cat. no. ABIN101961) and incubated on a nutator for 1 hour at room temperature. Beads were washed 3 times with 1 mL Dig-wash buffer, resuspended in 100 μL Dig-300 buffer containing 1:250 pA-Tn5 (diagenode, cat. no. C01070001) and incubated on a nutator for 1 hour at room temperature. Beads were washed and tagmentation was performed as described above for NanoTag. DNA was extracted with phenol-chloroform-isoamyl alcohol extraction as described in[32] and stored at –20^°^C. CUT&Tag libraries were amplified using 14 cycles and purified as described above for NanoTag.

### Sequencing

NanoTag and CUT&Tag libraries were serially diluted to equal molarity and pooled to 400 pM and 10% PhiX was added to the pool. Sequencing was performed on the Illumina NextSeq2000 using 50bp paired-end sequencing. Since NanoTag libraries show very high duplication rates, sequencing was performed aiming for 25 million read pairs per sample.

## Data availability

Datasets generated here were deposited to the European Nucleotide Archive under accession PRJEB77284. Public datasets used were: GSE208145[34] (CUT&RUN profiling CTCF in mESC CTCF-GFP-AID cell line (the same cell line used in this study)), GSE181104[35] (CUT&RUN profiling Nanog and CUT&Tag profiling H3K4me3 in mESC), GSE128907[25] (ChIP-seq data profiling TAF3 in mESC (the same cell line used in this study), GSE136488[36] (ChIP-seq data profiling Nanog and CTCF in mESC), GSE136479[36] (ChIP-seq data profiling H3K4me3 and H3K27me in mESC).

## Sequencing data analysis

Data was analyzed as described in[37]. Data obtained from cells expressing the GFP-tagged target was always compared to data from wild-type (WT) cells for both NanoTag and CUT&Tag. Briefly, fastq files were trimmed using trim_galore[38] and aligned to the mm10 genome using Bowtie2[39]. Unmapped, unpaired, low-quality, blacklisted, mitochondrial and duplicated reads were filtered using SAMtools[40]. Peaks were called using MACS2 using WT libraries as controls. Peaks called using downsampled CUT&Tag data were used for all downstream comparisons. Tracks were created using bamCompare with controls (--normalizeUsing BPM –-binSize 20). Heatmaps and correlation matrices were created using deeptools. All correlation matrices were generated using the alignment (bam) files. For CUT&Tag data, files were downsampled to match the sequencing depth of the NanoTag files (after filtering, Supp. Table 2). For ChIP-seq and CUT&RUN data, downsampling was not performed. Correlations were calculated at peaks common to the NanoTag, CUT&Tag and ChIP-seq datasets for the respective target. To generate coverage tracks for heatmaps, replicate alignment files were merged (and, for CUT&Tag downsampled to match the sequencing depth of NanoTag data). Heatmaps were generated using the coverage tracks signal around common peaks for each target. Coverage files were not background normalized, as in [37]. Peaks were annotated using ChIPseeker[41] and gene ontology (GO) analyses were performed using rGREAT[42]. The most enriched 14 GO terms common between NanoTag and CUT&Tag were displayed in the GO analysis plots.

## Library enrichment qPCR

To check NanoTag (and CUT&Tag) libraries for enrichment of fragments around desired target loci before sequencing, we devised a qPCR-based strategy. Primers were designed based on ENCODE ChIP-seq data for each target as follows: ENCODE peaks were ranked by enrichment score and two primer pairs were designed for each of the 10 strongest peaks per target. Primers were validated using gDNA extracted from the mES cell line used for each target. Primers for H3K4me3, Nanog and CTCF loci can be found below. Libraries were diluted 6-fold and 2 µL of diluted library was combined with 0.2 µL of 10 µM forward/reverse primer mix, 5 µL of SybrGreen qPCR master mix and 2.8 µL of water. qPCR was conducted using the following program: 95^°^C for 5 min and 50 cycles of: 95^°^C for 10 seconds, 60^°^C for 10 seconds, 72^°^C for 10 seconds. To determine the enrichment of NanoTag libraries at target loci, the difference in amplification latency between the libraries from cells expressing the GFP-tagged target and wild-type cells was compared at target loci and normalized to amplification at the gene desert.

## Enrichment qPCR primers

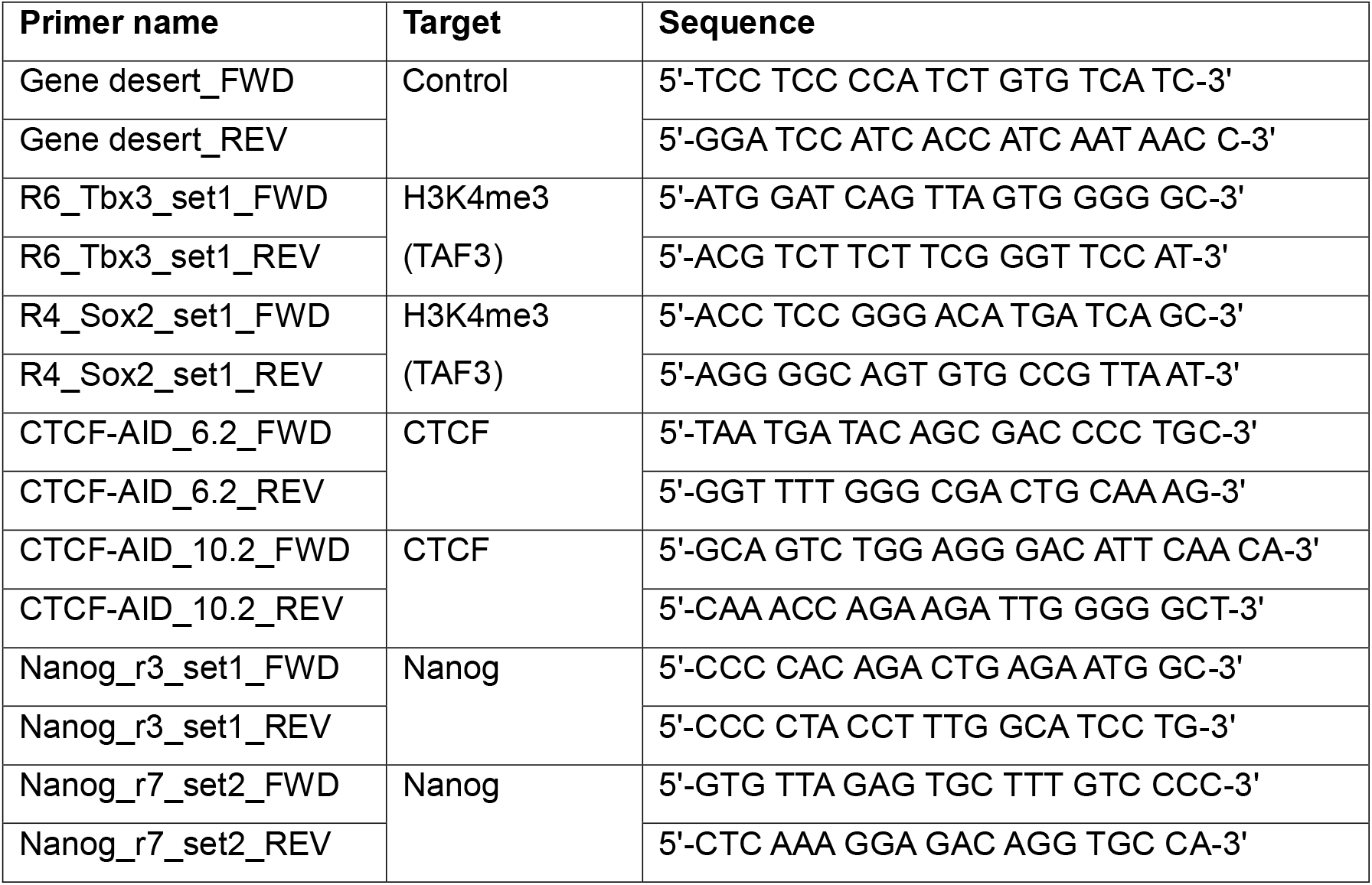

## Authors’ contributions

MAD developed the original idea, conceived and conducted experiments, analyzed data, produced graphs and figures, interpreted results and wrote the manuscript. LCS and RGAM helped conceive experiments and interpret results in consultation with PLG. IMM contributed to project conception and conduction, and data interpretation, provided guidance, corrected the manuscript and provided resources and funding. All authors contributed to the writing of the manuscript.

## Acknowledgements

We thank Martin Roszkowski and Rodrigo Villaseñor for their invaluable help with conceptualizing the NanoTag principle. We thank Tuncay Baubec, Timm Schroeder and Elphège Nora for providing us with mESC lines. We thank Alessandra Stürchler and Bogdan Mateescu for helpful discussions on experimental design and for critical feedback on the manuscript and Kerem Uzel for valuable help with the bioinformatic analyses. We thank Ellen Jaspers for her editorial help. We thank the Functional Genomics Center Zurich and the Protein Production and Structure Core Facility at EPFL for services and feedback. We thank all members of the Mansuy lab for valuable discussions. The Mansuy lab received funding from the University Zurich, ETH Zurich, the Swiss National Science Foundation (grant number 31003A_175742/1), ETH grants (ETH-10 15-2 and ETH-17 13-2), the National Centre of Competence in Research (NCCR) RNA&Disease funded by the Swiss National Science Foundation (grant number 182880/Phase 2 and 205601/Phase 3), the Hochschulmedizin Flagship Project “STRESS”, the European Union Horizon 2020 Research Innovation Program EarlyCause (Grant number 848158), the European Union projects FAMILY and HappyMums funded by the Swiss State Secretariat for Education, Research and Innovation (SERI), the FreeNovation grant from Novartis Forschungsstiftung and the Escher Family Fund. Rodrigo G. Arzate-Mejía received an ETH Postdoctoral Fellowship (grant number 20-1 FEL-28).

## Notes

### Competing Interest Statement

The authors have declared no competing interest.

## References

1. Klein DC, Hainer SJ. Genomic methods in profiling DNA accessibility and factor localization. Chromosome Res. 2020;28:69–85.

2. Kaya-Okur HS, Wu SJ, Codomo CA, Pledger ES, Bryson TD, Henikoff JG, Ahmad K, Henikoff S. CUT&Tag for efficient epigenomic profiling of small samples and single cells. Nat Commun. 2019;10:1930.

3. Carter B, Ku WL, Kang JY, Hu G, Perrie J, Tang Q, Zhao K. Mapping histone modifications in low cell number and single cells using antibody-guided chromatin tagmentation (ACT-seq). Nat Commun. 2019;10:3747.

4. Wang Q, Xiong H, Ai S, Yu X, Liu Y, Zhang J, He A. CoBATCH for high-throughput single-cell epigenomic profiling. Mol Cell. 2019;76:206–216.

5. Bartlett DA, Dileep V, Handa T, Ohkawa Y, Kimura H, Henikoff S, Gilbert DM. High-throughput single-cell epigenomic profiling by targeted insertion of promoters (Tip-seq). J Cell Biol. 2021;220:e202103078.

6. Gopalan S, Wang Y, Harper NW, Garber M, Fazzio TG. Simultaneous profiling of multiple chromatin proteins in the same cells. Mol Cell. 2021;81:4736–4746.

7. Meers MP, Llagas G, Janssens DH, Codomo CA, Henikoff S. Multifactorial profiling of epigenetic landscapes at single-cell resolution using MulTI-Tag. Nat Biotechnol. 2023;41:708–716.

8. Bartosovic M, Castelo-Branco G. Multimodal chromatin profiling using nanobody-based single-cell CUT&Tag. 2023;41:794–805.

9. Stuart T, Hao S, Zhang B, Mekerishvili L, Landau DA, Maniatis S, Satija R, Raimondi I. Nanobody-tethered transposition enables multifactorial chromatin profiling at single-cell resolution. Nat Biotechnol. 2023;41:806–812.

10. Janssens DH, Greene JE, Wu SJ, Codomo CA, Minot SS, Furlan SN, Ahmad K, Henikoff S. Scalable single-cell profiling of chromatin modifications with sciCUT&Tag. Nat Protoc. 2024;19:83–112.

11. Moleri P, Wilkins BJ. Unnatural amino acid crosslinking for increased spatiotemporal resolution of chromatin dynamics. Int J Mol Sci. 2023;24:12879.

12. Salma M, Andrieu-Soler C, Deleuze V, Soler E. High-throughput methods for the analysis of transcription factors and chromatin modifications: low input, single cell and spatial genomic technologies. Blood Cells Mol Dis. 2023;101:102745.

13. Partridge EC, Watkins TA, Mendenhall EM. Every transcription factor deserves its map: scaling up epitope tagging of proteins to bypass antibody problems. BioEssays. 2016;38:801–811.

14. Jin B, Odongo S, Radwanska M, Magez S. Nanobodies: a review of generation, diagnostics and therapeutics. Int J Mol Sci. 2023;24:5994.

15. Van Ingen H, Van Schaik FMA, Wienk H, Ballering J, Rehmann H, Dechesne AC, Kruijzer JA, Liskamp RM, Timmers HT, Boelens R. Structural insight into the recognition of the H3K4me3 mark by the TFIID subunit TAF3. Structure. 2008;16:1245–1256.

16. Bernstein BE, Mikkelsen TS, Xie X, Kamal M, Huebert DJ, Cuff J, Fry B, Meissner A, Wernig M, Plath K, Jaenisch R, Wagschal A, Feil R, Schreiber SL, Lander ES. A bivalent chromatin structure marks key developmental genes in embryonic stem cells. Cell. 2006;125:315–326.

17. Bernstein BE, Kamal M, Lindblad-Toh K, Bekiranov S, Bailey DK, Huebert DJ, McMahon S, Karlsson EK, Kulbokas EJ, Gingeras TR, Schreiber SL, Lander ES. Genomic maps and comparative analysis of histone modifications in human and mouse. Cell. 2005;120:169–181.

18. Wang M, Zhang Y. Tn5 transposase-based epigenomic profiling methods are prone to open chromatin bias. 2021 Jul. bioRxiv 2021.07.09.451758.

19. Filipczyk A, Marr C, Hastreiter S, Feigelman J, Schwarzfischer M, Hoppe PS, Loeffler D, Kokkaliaris KD, Endele M, Schauberger B, Hilsenbeck O, Skylaki S, Hasenauer J, Anastassiadis K, Theis FJ, Schroeder T. Network plasticity of pluripotency transcription factors in embryonic stem cells. Nat Cell Biol. 2015; 17:1235–1246.

20. Nora EP, Goloborodko A, Valton AL, Gibcus JH, Uebersohn A, Abdennur N, Dekker N, Mirny LA, Bruneau BG. Targeted degradation of CTCF decouples local insulation of chromosome domains from genomic compartmentalization. Cell. 2017;169:930–944.

21. Nordin A, Pagella P, Zambanini G, Cantu C. Exhaustive identification of genome-wide binding events of transcriptional regulators, Nucleic Acids Research, 2024;52:7.

22. Hu D, Abbasova L, Schilder BM, Nott A, Skene NG, Marzi SJ. CUT&Tag recovers up to half of ENCODE ChIP-seq peaks. bioRxiv preprint. 2022.03.30.486382.

23. McCombie WR, McPherson JD. Future promises and concerns of ubiquitous next-generation sequencing. Cold Spring Harb Perspect Med. 2019;9:a025783.

24. Monteiro CJ, Heery DM, Whitchurch JB. Modern approaches to mouse genome editing using the CRISPR-Cas toolbox and their applications in functional genomics and translational research. In: Passos GA, editor. Genome Ed Biomed Sci 2023:13–40.

25. Villaseñor R, Pfaendler R, Ambrosi C, Butz S, Giuliani S, Bryan E, Sheahan TW, Gable AL, Schmolka N, Manzo M, Wirz J, Feller C, von Mering C, Aebersold R, Voigt P, Baubec T. ChromID identifies the protein interactome at chromatin marks. Nat Biotechnol. 2020;38:728–736.

26. Scholz O, Hansen S, Plückthun A. G-quadruplexes are specifically recognized and distinguished by selected designed ankyrin repeat proteins. Nucleic Acids Res. 2014;42:9182–9194.

27. Hayashi-Takanaka Y, Yamagata K, Wakayama T, Stasevich TJ, Kainuma T, Tsurimoto T, Tachibana M, Shinkai Y, Kurumizaka H, Nozaki N, Kimura H. Tracking epigenetic histone modifications in single cells using Fab-based live endogenous modification labeling. Nucleic Acids Res. 2011;39:6475–6488.

28. Moeglin E, Desplancq D, Stoessel A, Massute C, Ranniger J, McEwen AG, Zeder-Lutz G, Oulad-Abdelghani M, Chiper M, Lafaye P, Di Ventura B, Didier P, Poterszman A, Weiss E. A Novel Nanobody Precisely Visualizes Phosphorylated Histone H2AX in Living Cancer Cells under Drug-Induced Replication Stress. Cancers. 2021;13:3317.

29. Nguyen-Duc T, Peeters E, Muyldermans S, Charlier D, Hassanzadeh-Ghassabeh G. Nanobody-based chromatin immunoprecipitation/micro-array analysis for genome-wide identification of transcription factor DNA binding sites. Nucleic Acids Res. 2013;41:e59.

30. Katoh Y, Nozaki S, Hartanto D, Miyano R, Nakayama K. Architectures of multisubunit complexes revealed by a visible immunoprecipitation assay using fluorescent fusion proteins. J Cell Sci. 2015;128:2351–2362.

31. Chen W, Gardeux V, Meireles-Filho A, Deplancke B. Profiling of single-cell transcriptomes. Curr Protoc Mouse Biol. 2017;7:145–175.

32. S Kaya-Okur H. Bench top CUT&Tag v3. protocols.io. 2020. https://www.protocols.io/view/bench-top-cut-amp-tag-bcuhiwt6.

33. Grandi FC, Modi H, Kampman L, Corces MR. Chromatin accessibility profiling by ATAC-seq. Nat Protoc. 2022;17:1518–1552.

34. Wulfridge P, Yan Q, Rell N, Doherty J, Jacobson S, Offley S, Deliard S, Feng K, Phillips-Cremins J, Gardini A, Sarma K. G-quadruplexes associated with R-loops promote CTCF binding. Mol Cell. 2023;83:3064–3079.

35. Thompson JJ, Lee DJ, Mitra A, Frail S, Dale RK, Rocha PP. Extensive co-binding and rapid redistribution of NANOG and GATA6 during emergence of divergent lineages. Nat Commun. 2022;13:4257.

36. The ENCODE Project Consortium. An integrated encyclopedia of DNA elements in the human genome. Nature. 2012;489:57–74.

37. Zheng Y, Ahmad K, Henikoff S. CUT&Tag data processing and analysis tutorial v1. Github. 2020. https://yezhengstat.github.io/CUTTag_tutorial/.

38. Krueger F, James F, Ewels P, Afyounian E, Weinstein M, Schuster-Boeckler B. FelixKrueger/TrimGalore: v0.6.7 – DOI via Zenodo (0.6.7). Zenodo. 10.5281/zenodo.5127899 (2021).

39. Langmead B, Salzberg SL. Fast gapped-read alignment with Bowtie 2. Nat Methods. 2012;9:357–359.

40. Danecek P, Bonfield JK, Liddle J, Marshall J, Ohan V, Pollard MO, Whitwham A, Keane T, McCarthy SA, Davies RM, Li H. Twelve years of SAMtools and BCFtools. GigaScience. 2021;10:giab008.

41. Yu G, Wang L-G, He Q-Y. ChIPseeker: an R/Bioconductor package for ChIP peak annotation, comparison and visualization. Bioinformatics. 2015;31:2382–2383.

42. Gu Z, Hübschmann D. rGREAT: an R/bioconductor package for functional enrichment on genomic regions. Bioinformatics. 2023;39:btac745.

43. Huang H, Zhu Q, Jussila A, Han Y, Bintu B, Kern C, et al. CTCF mediates dosage– and sequence-context-dependent transcriptional insulation by forming local chromatin domains. Nat Genet. 2021;53:1064–1074.

