## Supplemental information for "NanoTag - an IgG-free method for mapping DNA-protein interactions"

**Supplemental Figure 1 Dimitriu et al**

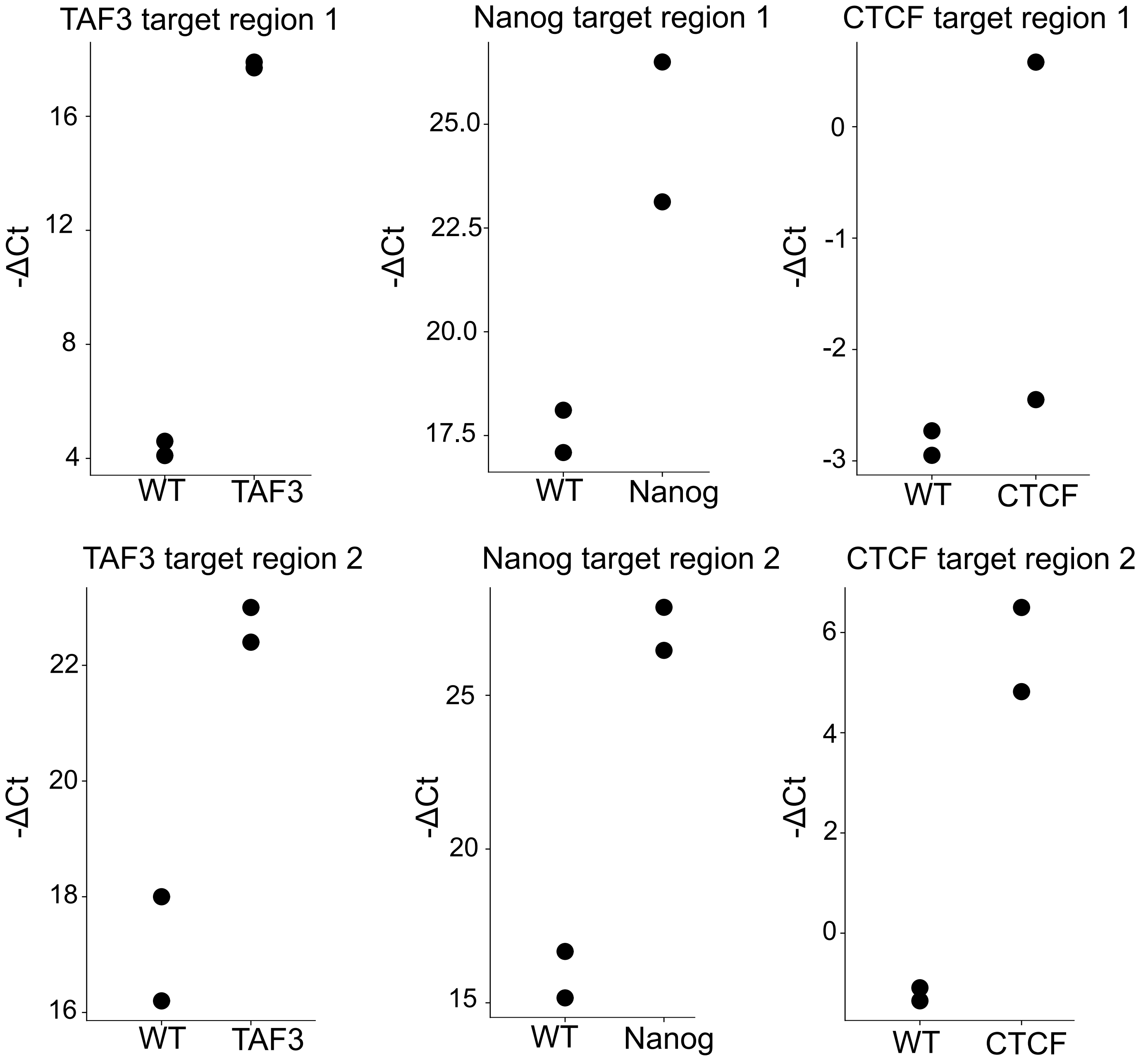

Supplemental Figure 2 Dimitriu et al

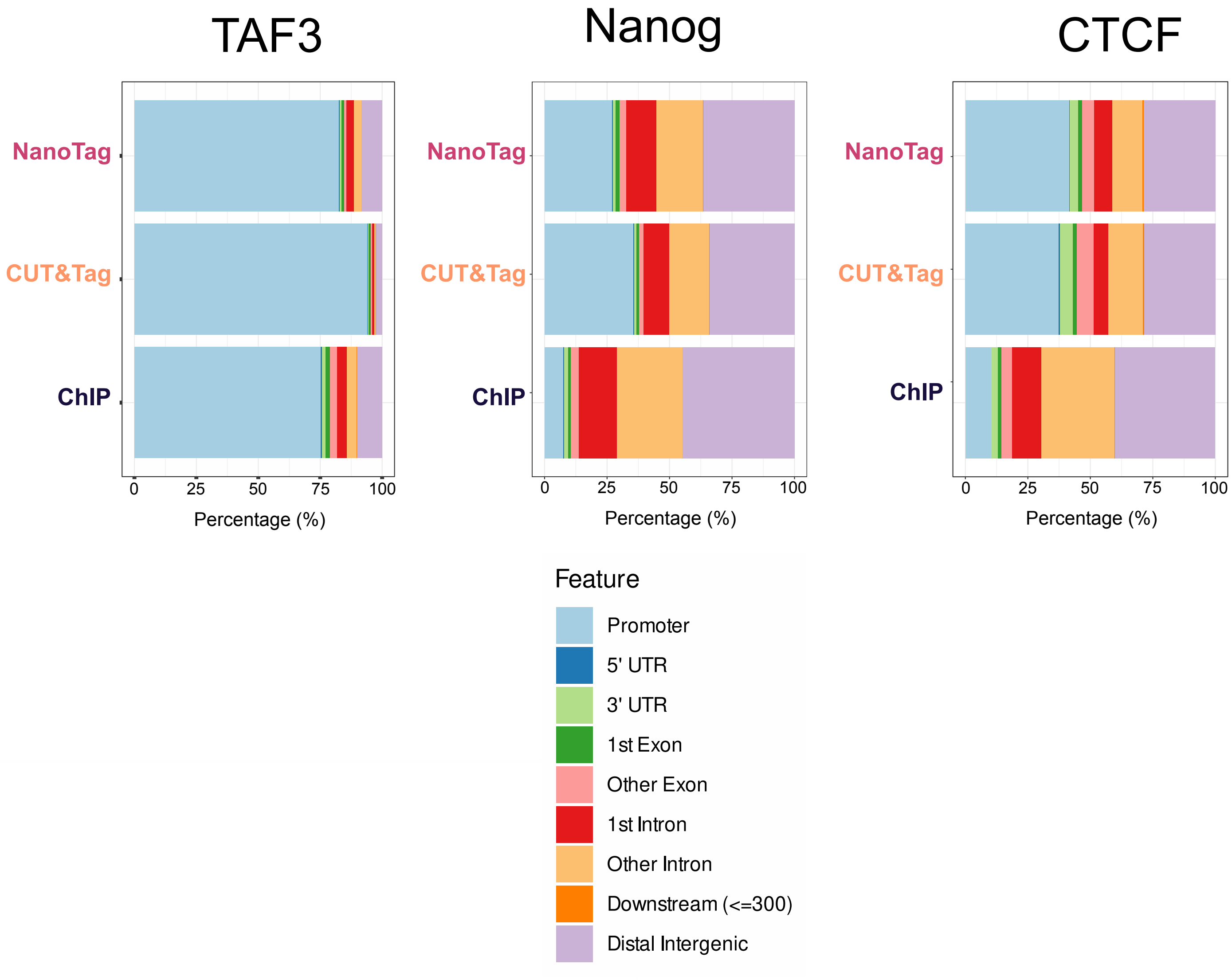

### Supplemental figure 3 Dimitriu et al

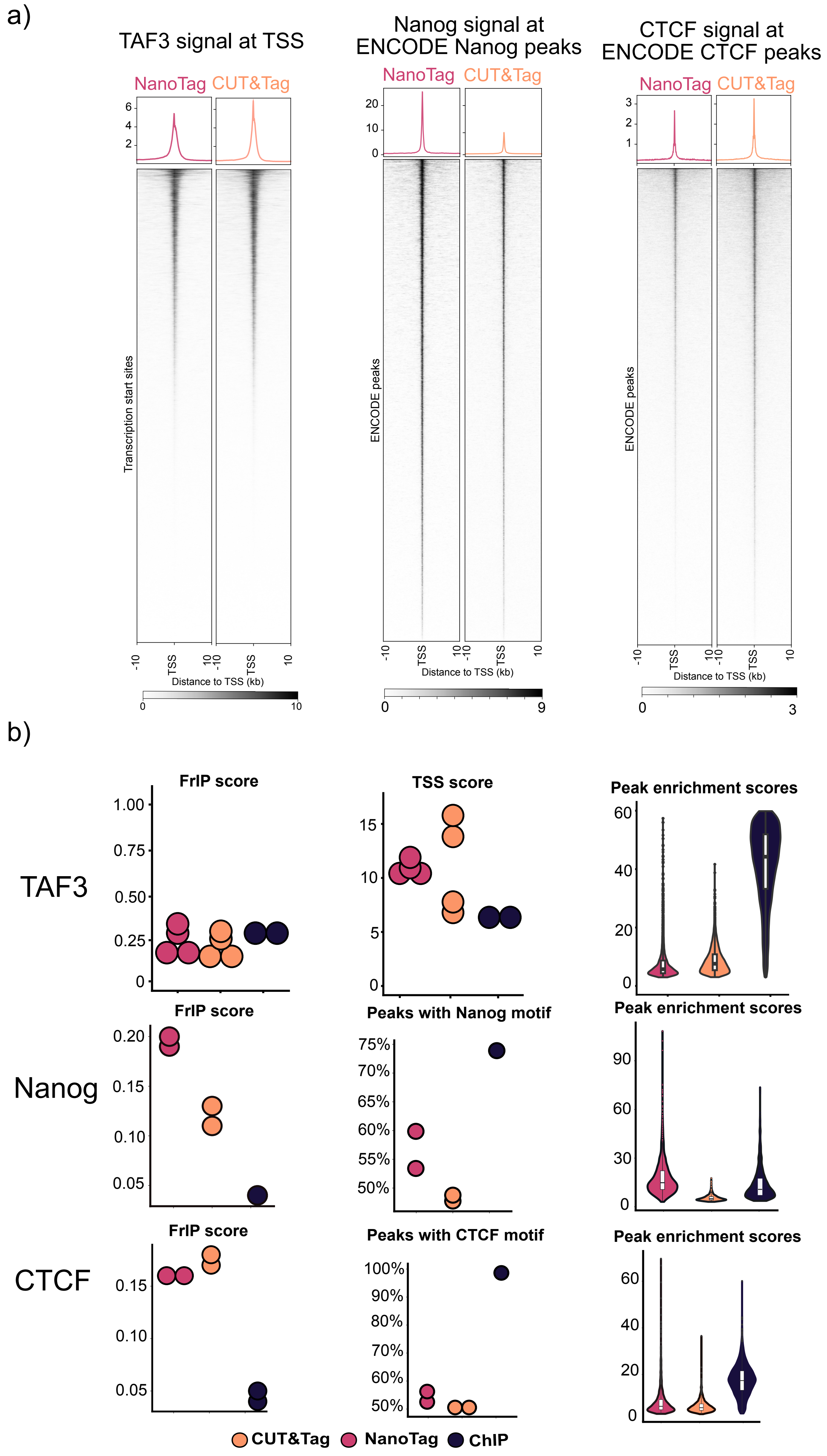

**Supplemental Figure 4 Dimitriu et al**

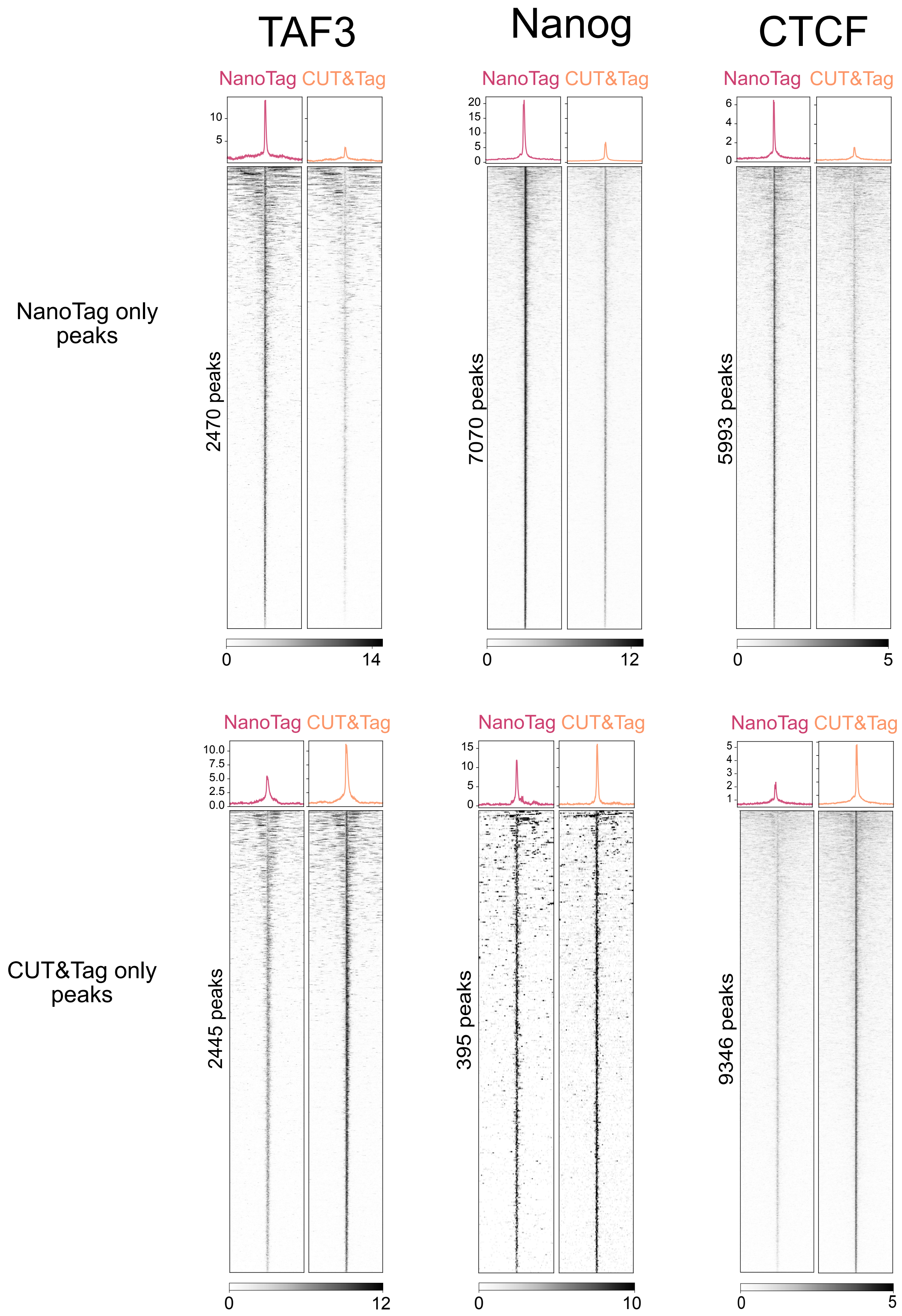

### Supplemental Figure 5 Dimitriu et al

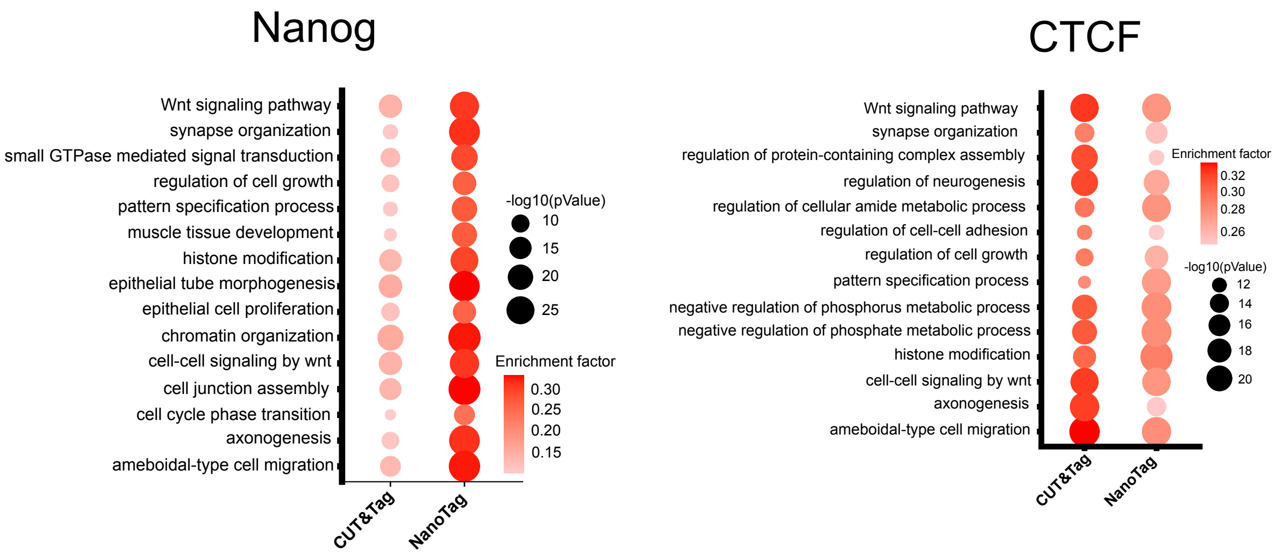

**Supplemental Figure 6 Dimitriu et al**

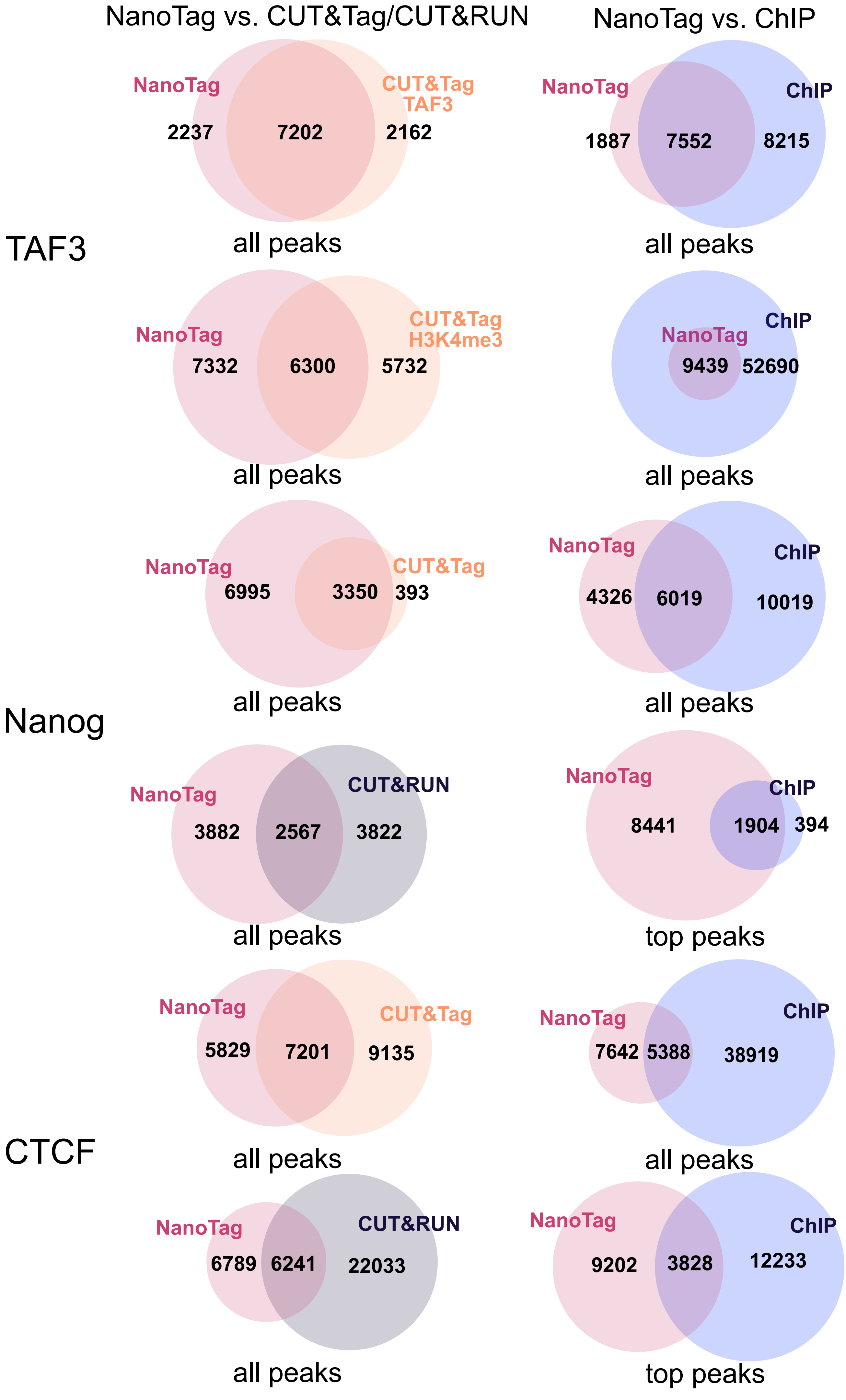

### Supplemental Table 1 Dimitriu et al

| Sample | Method | Cell line | Target | Replicate | Total_reads | Trimmed_reads | Filtered_reads | Mapping_rates | Duplication_rates | Peaks |
|---|---|---|---|---|---|---|---|---|---|---|
| NanoTag_mESC_TAF-GFP_r1 | NanoTag | TAF3-GFP mESC | TAF3 | 1 | 9.36E+07 | 9.31E+07 | 1.89E+06 | 96.2% | 96% | 7747 |
| NanoTag_mESC_TAF-GFP_r1 | NanoTag | TAF3-GFP mESC | TAF3 | 2 | 9.97E+07 | 9.94E+07 | 2.40E+06 | 96.2% | 95% | 8067 |
| NanoTag_mESC_Nanog-GFP_r1 | NanoTag | Nanog-Venus mESC | Nanog | 1 | 1.52E+08 | 1.49E+08 | 1.92E+06 | 97.2% | 97% | 8452 |
| NanoTag_mESC_Nanog-GFP_r2 | NanoTag | Nanog-Venus mESC | Nanog | 2 | 1.06E+08 | 1.05E+08 | 3.54E+06 | 97.2% | 95% | 8343 |
| NanoTag_mESC_CTCF-GFP_r1 | NanoTag | CTCF-GFP mESC | CTCF | 1 | 1.53E+08 | 1.42E+08 | 1.12E+06 | 96.3% | 98% | 9869 |
| NanoTag_mESC_CTCF-GFP_r2 | NanoTag | CTCF-GFP mESC | CTCF | 2 | 1.32E+08 | 1.25E+08 | 9.99E+05 | 96.3% | 98% | 8626 |
| NanoTag_mESC_WT | NanoTag | WT mESC | NA | 1 | 3.23E+07 | 3.14E+07 | 4.06E+05 | 95.0% | 97% | NA |
| CUT&Tag_mESC_TAF-GFP_r1 | CUT&Tag | TAF3-GFP mESC | TAF3 | 1 | 8.41E+07 | 8.41E+07 | 3.50E+07 | 98.1% | 50% | 29513 |
| CUT&Tag_mESC_TAF-GFP_r2 | CUT&Tag | TAF3-GFP mESC | TAF3 | 2 | 1.08E+08 | 1.08E+08 | 4.16E+07 | 98.2% | 54% | 32469 |
| CUT&Tag_mESC_Nanog-GFP_r1 | CUT&Tag | Nanog-Venus mESC | Nanog | 1 | 1.05E+08 | 1.05E+08 | 3.20E+07 | 98.1% | 60% | 66425 |
| CUT&Tag_mESC_Nanog-GFP_r2 | CUT&Tag | Nanog-Venus mESC | Nanog | 2 | 1.16E+08 | 1.16E+08 | 3.84E+07 | 98.0% | 57% | 65995 |
| CUT&Tag_mESC_CTCF-GFP_r1 | CUT&Tag | CTCF-GFP mESC | CTCF | 1 | 9.92E+07 | 9.91E+07 | 4.28E+07 | 98.0% | 48% | 81512 |
| CUT&Tag_mESC_CTCF-GFP_r2 | CUT&Tag | CTCF-GFP mESC | CTCF | 2 | 1.08E+08 | 1.07E+08 | 4.42E+07 | 97.9% | 51% | 81830 |
| CUT&Tag_mESC_IgG | CUT&Tag | TAF3-GFP mESC | NA | 1 | 1.18E+08 | 1.18E+08 | 7.98E+06 | 97.5% | 88% | NA |
| CUT&Tag_mESC_WT | CUT&Tag | WT mESC | NA | 2 | 1.06E+08 | 1.06E+08 | 1.37E+07 | 98.1% | 80% | NA |

### Supplemental Table 2 Dimitriu et al

| Sample | Method | Cell line | Target | Replicate | Filtered_reads | Peaks | Reproducible | % of total |
|---|---|---|---|---|---|---|---|---|
| NanoTag_mESC_TAF-GFP_r1 | NanoTag | TAF3-GFP mESC | TAF3 | 1 | 1.89E+06 | 7747 | 6374 | 82.3 |
| NanoTag_mESC_TAF-GFP_r1 | NanoTag | TAF3-GFP mESC | TAF3 | 2 | 2.40E+06 | 8067 | | 79.0 |
| NanoTag_mESC_Nanog-GFP_r1 | NanoTag | Nanog-Venus mESC | Nanog | 1 | 1.92E+06 | 8452 | 6449 | 76.3 |
| NanoTag_mESC_Nanog-GFP_r2 | NanoTag | Nanog-Venus mESC | Nanog | 2 | 3.54E+06 | 8343 | | 77.3 |
| NanoTag_mESC_CTCF-GFP_r1 | NanoTag | CTCF-GFP mESC | CTCF | 1 | 1.12E+06 | 9869 | 5465 | 55.4 |
| NanoTag_mESC_CTCF-GFP_r2 | NanoTag | CTCF-GFP mESC | CTCF | 2 | 9.99E+05 | 8626 | | 63.4 |
| CUT&Tag_mESC_TAF-GFP_r1 | CUT&Tag | TAF3-GFP mESC | TAF3 | 1 | 2.13E+06 | 7862 | 7397 | 94.1 |
| CUT&Tag_mESC_TAF-GFP_r2 | CUT&Tag | TAF3-GFP mESC | TAF3 | 2 | 2.12E+06 | 8898 | | 83.1 |
| CUT&Tag_mESC_Nanog-GFP_r1 | CUT&Tag | Nanog-Venus mESC | Nanog | 1 | 2.72E+06 | 2597 | 1984 | 76.4 |
| CUT&Tag_mESC_Nanog-GFP_r2 | CUT&Tag | Nanog-Venus mESC | Nanog | 2 | 2.72E+06 | 3129 | | 63.4 |
| CUT&Tag_mESC_CTCF-GFP_r1 | CUT&Tag | CTCF-GFP mESC | CTCF | 1 | 1.03E+06 | 11092 | 7161 | 64.6 |
| CUT&Tag_mESC_CTCF-GFP_r2 | CUT&Tag | CTCF-GFP mESC | CTCF | 2 | 1.02E+06 | 12406 | | 57.7 |

### Supplemental Figure 7 Dimitriu et al

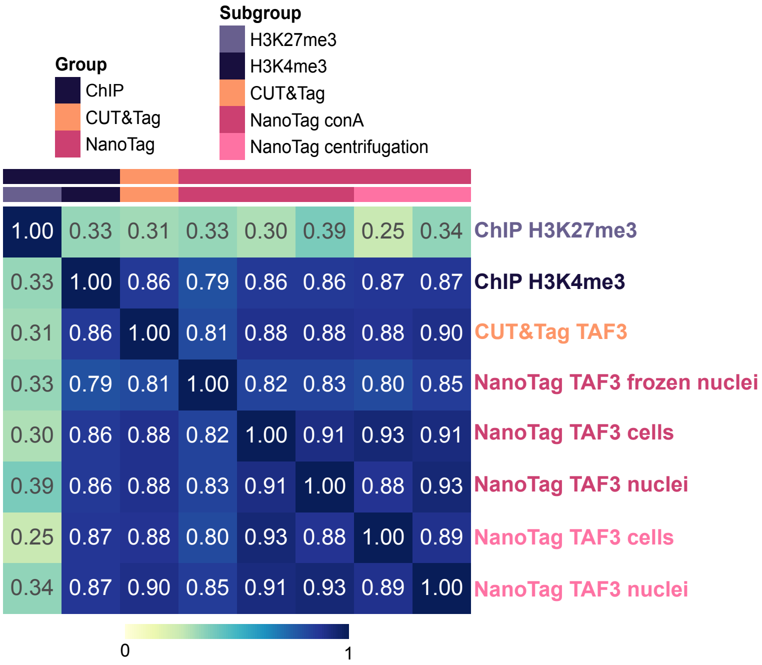

**Supplemental Figure 1** NanoTag libraries are enriched for fragments containing expected target binding sites. For each target profiled using NanoTag, libraries were amplified with primers targeting two different regions where the protein is expected to bind. Numbers of cycles required for amplification were compared for each target between libraries prepared from cells expressing the GFP-tagged target and libraries prepared from control (WT) cells. ΔCt value of each sample represents the amplification at the target region normalized to amplification at the gene desert control region. -ΔCt values were plotted to show the higher amplification of libraries derived from GFP-expressing cells compared to WT cells.

**Supplemental Figure 2** Summary of NanoTag, CUT&Tag and ChIP-seq peaks distribution over different genomic elements for each target. For each method, the peak set used represents peaks common to both replicates. NanoTag and CUT&Tag data were generated in-house, ChIP-seq peaks are from Villaseñor et al. for TAF3 and from ENCODE project for Nanog and CTCF.

**Supplemental Figure 3** Enrichment of NanoTag and CUT&Tag data and comparisons to ChIP-seq data for each target. a) Profile plots and heatmaps of NanoTag and CUT&Tag data at TSS when profiling TAF3 and at ENCODE peaks when profiling TF. b) Plots summarizing the FrIP scores at ENCODE peaks, the TSS scores (for TAF3), the proportion of peaks containing the expected TF motif (for Nanog and CTCF) and the enrichment score distribution for common peaks for NanoTag (magenta), CUT&Tag (orange) and ChIP-seq (dark blue) data. FrIP scores and proportion of peaks containing the TF motif were calculated individually for peak sets called in each replicate. Coverage of individual replicates was merged to calculate the enrichment score at common peaks. Motifs were identified in the 500 most significant peaks across the two replicates.

**Supplemental Figure 4** Profile plots and heatmaps showing NanoTag and CUT&Tag coverage at peaks called only in the NanoTag (top) or only in the CUT&Tag data (bottom) for each target.

**Supplemental Figure 5** Gene ontology analysis for NanoTag and CUT&Tag peaks when profiling Nanog and CTCF. The top 14 common GO terms are shown for each target with the corresponding enrichment score and p values.

**Supplemental Figure 6** Overlaps between NanoTag peak sets and CUT&Tag or CUT&Run (left) and between NanoTag peaks sets and ChIP-seq peak sets (right) for each target. Peaks were called by downsampling the CUT&Tag data to match the sequencing depth of NanoTag data (but without downsampling the CUT&Run and ChIP-seq data). For ChIP-seq data, comparison was done to all ChIP-seq peaks and to the top 15% most significant peaks separately.

**Supplemental Table 1.** Summary of NanoTag and CUT&Tag raw and processed data, including alignment rates, duplication rates and peaks called for each replicate.

**Supplemental Table 2.** Summary of processed data used for all analyses. CUT&Tag files were downsampled to the average sequencing depth of the two NanoTag replicates for each target and peaks called on the downsampled dataset were used for further analyses. Reproducible peaks are peaks called in both replicates.

**Supplemental Figure 7** NanoTag protocol variations produce similar results with different input materials. Correlation matrix comparing coverage of NanoTag TAF3 data obtained using different inputs and different protocol variations (shades of pink) to CUT&Tag (orange) and ChIP-seq (purple) data. Read-level Spearman correlation of TAF3-NanoTag samples prepared using different protocol versions and input material to TAF3-CUT&Tag and ChIP-seq. NanoTag was performed using conA beads as for CUT&Tag (NanoTag conA, dark pink) on fresh cells, fresh nuclei or frozen nuclei and using centrifugation (light pink, see Methods for details) to pellet cells or nuclei. NanoTag and CUT&Tag data was produced in-house, ChIP-seq data is from ENCODE.
